# Pathway-based polygenic risk implicates GO: 17144 drug metabolism in recurrent depressive disorder

**DOI:** 10.1101/209544

**Authors:** Anna R. Docherty, Arden Moscati, T. Bernard Bigdeli, Alexis K. Edwards, Roseann Peterson, Fuzhong Yang, Daniel E. Adkins, John S. Anderson, Jonathan Flint, Kenneth S. Kendler, Silviu-Alin Bacanu

## Abstract

The Psychiatric Genomics Consortium (PGC) has made major advances in the molecular etiology of MDD, confirming that MDD is highly polygenic, with any top risk loci conferring a very small proportion of variance in case-control status (1). Pathway enrichment results from PGC meta-analyses can also be used to help inform molecular drug targets. Prior to any knowledge of molecular biomarkers for MDD, drugs targeting molecular pathways have proved successful in treating MDD. However, it is possible that with information from PGC analyses, examining specific molecular pathway(s) implicated in MDD can further inform our study of molecular drug targets. Using a large case-control GWAS based on low-coverage whole genome sequencing (N = 10,640), we derived polygenic risk scores for MDD and for MDD specific to each of over 300 molecular pathways. We then used these data to identify sets of scores significantly predictive of case status, accounting for critical covariates. Over and above global polygenic risk for MDD, polygenic risk within the GO: 17144 drug metabolism pathway significantly predicted recurrent depression. In transcriptomic analyses, two pathway genes yielded suggestive signals at FDR *q*-values = .054: *CYP2C19* (family of Cytochrome P450) and *CBR1* (Carbonyl Reductase 1). Because the neuroleptic carbamazepine is a known inducer of *CYP2C19*, future research might examine whether drug metabolism PRS has any influence on clinical presentation and treatment response. Overall, results indicate that pathway-based risk might inform treatment of severe depression. We discuss limitations to the generalizability of these preliminary findings, and urge replication in future research.

## Introduction

Recurrent major depressive disorder (MDD) is a complex phenotype with limited established associations with biological markers and molecular (gene) pathways. The China, Oxford and Virginia Commonwealth University Experimental Research on Genetic Epidemiology (CONVERGE) study and the Psychiatric Genomics Consortium (PGC) have made significant advances in the molecular etiology of MDD, confirming that MDD is highly polygenic, with aggregation of top loci effects accounting for a small proportion of the variance in case-control status (1,2).

With these genomic data, it is increasingly feasible to begin to isolate and examine molecular drug targets for MDD. One way to work toward this aim is by inspecting molecular pathway enrichment in relation to severe MDD diagnosis, or in relation to symptom presentation in severe cases. Though, it is important to present caveats: molecular pathway analyses are fraught with potential for spurious findings and require 1) large sample sizes, 2) strict correction for multiple comparison, 3) a grain of salt, given the diffuse and quite polygenic signal from even very large GWAS meta-analyses, and 4) replication. It is also important to note that available information about molecular pathways is far from complete.

Keeping these important points in mind, we also know that many molecular pathways (MPs) are enriched for MDD signal, and that neuroleptics targeting specific neurotransmitter pathways are efficacious. Thus, it is possible that MPs have some influence on the presentation of and risk for severe MDD, and potentially on drug response. Given the aims of genomic research to identify promising drug targets for MDD, the identification of potential molecular classifiers of drug response is highly desirable.

To this aim, we examined global polygenic risk for MDD, and also polygenic risk for MDD specific to molecular pathways, in the presentation of severe MDD in a large case-control sample of Han Chinese descent (*N* = 10,064; CONVERGE) (3). CONVERGE aimed at identifying genetic risk factors for recurrent MDD among a rigorously ascertained cohort, and presents several advantages for genetic study. The sample is entirely female, thus reducing heterogeneity from sex differences. Further, unlike most studies of its size, CONVERGE subjects all have 4 Han Chinese grandparents, thus reducing noise stemming from population stratification. Finally, all subjects were ascertained for recurrent depression, rather than a single episode, to increase severity and thus potential genetic signal. This ascertainment method, and the size of the study, allowed CONVERGE to identify the first risk loci for MDD.

Our analysis used summary statistics from a leave-one-out GWAS of MDD within CONVERGE. We constructed Bayesian subject-level genome-wide MDD PRS and MDD PRS specific to pruned molecular pathways. Scores and covariates were regressed onto case status, and pathway scores with significant signal above and beyond global MDD polygenic risk were then examined with within-pathway, gene-based PRS.

## Methods and Materials

### Sampling

Recurrent MDD cases and healthy controls were recruited from 51 mental health centers and psychiatric departments of general medical hospitals across 21 provinces of China. Please refer to previously published research for full details of sample ascertainment (3). We controlled for potential clinical heterogeneity by recruiting only female participants, and to control for ethnic stratification, only participants whose grandparents (all four) were of Han Chinese descent were recruited to participate. Cases and controls (age *M* (*SD*) = 44.4 (8.9) and 47.7 (5.6), respectively) were excluded for diagnosis of bipolar disorder, any psychosis, and any diagnosis of intellectual disability. Cases had at least two major depressive episodes with the first episode occurring prior to age 50, and could not have abused drugs or alcohol prior to their first episode of depression. Controls were clinically screened to rule out prior depressive episodes and had to be at least 40 years of age, past the age window of typical MDD onset. The study protocol was approved centrally by the Ethical Review Board of Oxford University, and by the ethics committees of all of the participating hospitals in China. All participants provided written informed consent.

### DNA Sequencing

DNA extraction, sequencing, genotyping, and imputation details have been reported (4). Briefly, DNA was extracted from saliva using Oragene and sequenced reads were obtained from Illumina Hiseq machines aligned to Genome Reference Consortium Human Build 37 patch release 5 (GRCh37.p5) with Stampy (v1.0.17) with default parameters (5). Reads consisting of base quality <= 5 or containing adaptor sequencing were filtered out. Alignments were indexed in BAM format^5^ using Samtools (v0.1.18) (6) and PCR duplicates were marked for downstream filtering using Picardtools (v1.62). The Genome Analysis Toolkit’s (GATK, version 2.6) (7) BaseRecalibrator created recalibration tables to screen known SNPs and INDELs in the BAM files from dbSNP (version 137, excluding all sites added after version 129). GATKlite (v2.2.15) was used for subsequent base quality recalibration (BQSR) and removal of read pairs with improperly aligned segments as determined by Stampy.

### Calling and Imputation of Genotypes

GATK’s UnifiedGenotyper VariantRecalibrator (version 2.7-2-g6bda569) was used on post-BSQR files for variant discovery and genotyping at all polymorphic SNPs in 1000G Phase1 ASN panel (8) as well as variant quality score recalibration. A sensitivity threshold of 90% was applied for imputation after optimizing for Transition to Transversion ratios. Genotype likelihoods were calculated using a binomial mixture model implemented in SNPtools (version 1.0) (9) and imputation was performed at sites with no reference panel using BEAGLE (version 3.3.2) (10). A second round of imputation was performed at biallelic polymorphic SNPs using 1000G Phase 1 ASN haplotypes as a reference panel. To determine the final number of SNPs, we applied a conservative inclusion threshold for SNPs: 1) a *p*-value for violation *HWE* > 10-6, 2) information score > 0.9, and 3) minor allele frequency in CONVERGE > 0.5%.

### Diagnostic Assessments

Participant interview sessions lasted approximately two hours. Interviewers were largely trained psychiatrists with a small number representing post-graduate medical students or psychiatric nurses, and all were clinically trained by the CONVERGE team for at least one week. Interviews were recorded and included an assessment of psychopathology, demographic characteristics, and psychosocial functioning. Trained editors listened to a portion of the interviews in order to provide ratings of interview quality. We excluded participants who had incomplete assessment information or were lacking high-quality genetic data, to arrive at a final 4,728 controls and 5,612 case samples for analysis.

### Polygenic Risk for MDD and Pathway-Based Risk as a Predictor of Case Status

PRS were constructed for global MDD and for MDD within each of the molecular pathways, using leave-one-out GWAS analyses of the CONVERGE sample. A list of pathways was derived from independent, previously published pathway enrichment statistics from the PGC (11). Gene lists corresponding to each pathway were extracted from the relevant gene ontology database, with the number of genes per pathway ranging from 8 to 598 (mean = 9). Python-based LDpred (12), a polygenic risk scoring software, allows for the modeling of LD to weight the relative contributions of syntenic variants to the outcome phenotype. LDpred uses postulated proportions of causal variants in the genome as Bayesian prior probabilities for PRS calculations, and a range of 8 different priors were used (proportions of 1, 0.3, 0.1, 0.03, 0.01, 0.003, and 0.001) to construct scores. All pathway scores were LD pruned and merged based on excessive correlation between scores (R^2^< 0.5). Stepwise regressions were run with all PRS variables and critical covariates as determined by Cai et al. (3). Model 1 included all risk scores and the two primary ancestry principal components. Model 2 included all risk scores, primary ancestry principal components, and the presence/absence of childhood adversity (a significant moderator of the genetic signal in previous CONVERGE GWAS).

## Results

Stepwise regression statistics for the best fitting model are presented in Table 1, each parameter in the model remaining significant after FDR correction. The best-fitting stepwise regression included three robust predictors: Global MDD, one ancestry PC, and the MDD PRS for the GO:17144 drug metabolism pathway. Model 2 results are presented in Table 2. Global MDD risk and GO:17144 pathway-based risk remained significant in Model 2, where lifetime adversity (y/n) was entered as a covariate (and was found to be the most robust predictor). Of note, the stepwise models captured no increase in pathway-based risk by genome-wide MDD PRS.

**Table 1.** Best fitting stepwise regression model predicting MDD case status from global MDD PRS, molecular pathway-based PRS, and ancestry.

|  | Estimate | Standard Error | t-value | p-value |
| --- | --- | --- | --- | --- |
| Ancestry Principal Component 2 | -3.176 | 0.565 | -5.624 | $1.86 \times 10^{-8}$ |
| MDD global L1O PRS | -0.038 | 0.003 | -13.177 | $1.19 \times 10^{-39}$ |
| GO:17144 Drug Metabolism L1O PRS | -0.056 | 0.012 | -4.828 | $1.38 \times 10^{-6}$ |

**Table 2.** Best fitting stepwise regression model accounting for childhood adversity.

|  | Estimate | Standard Error | t-value | p-value |
| --- | --- | --- | --- | --- |
| Lifetime Adversity | 0.176 | 0.011 | 15.843 | $1.57 \times 10^{-56}$ |
| MDD global L1O PRS | -0.031 | 0.003 | -11.855 | $2.04 \times 10^{-32}$ |
| Ancestry Principal Component 2 | -3.989 | 0.503 | -7.933 | $2.13 \times 10^{-15}$ |
| GO:17144 Drug Metabolism L1O PRS | -0.050 | 0.010 | -4.845 | $1.27 \times 10^{-6}$ |

Follow-up gene-based analysis included MDD PRS calculation specific to the nine genes involved in the GO:17144 pathway. Regression onto case status resulted in 3 genes with suggestive signals (Table 3). Two of the three suggestive genes are influenced by the common neuroleptics carbamazepine and haloperidol. The first drug is an anti-convulsant commonly used for treating bipolar disorder and depression, and the latter an antipsychotic commonly used for treating acute psychosis. *CYP2C19* and *CBR1* both retained FDR q-values of .05.

**Table 3.** Suggestive genes in drug metabolism pathway after false discovery rate correction.

| Suggestive Genes in GO:17144 | Chr:Mbp start-end | FDR | Targets/antagonists |
| --- | --- | --- | --- |
| CYP2C19 (Cytochrome P450 Family 2 Subfamily C Member 19) | 10:94.763-10:94.853 | 0.05 | Carbamazepine |
| CBR1 (Carbonyl Reductase 1) | 21:36.070-21:36.073 | 0.05 | Haloperidol |
| FMO2 (Flavin Containing Monooxygenase 2) | 1:171.185-1:171.213 | 0.09 |  |

## Discussion

In the CONVERGE sample, MDD polygenic risk specific to GO:17144 drug metabolism pathway significantly predicted MDD case status. This effect remained significant while accounting for global MDD polygenic risk and for a genetic risk moderator, childhood adversity. These scores were based on MDD GWAS weights from leave-one-out analyses of the CONVERGE sample, so were entirely within-ancestry. This is the first study to our knowledge to test MDD PRS specific to molecular pathways in relation to recurrent MDD, and preliminary results suggest that GO:17144 may be relevant to clinical presentation.

Because MDD is genetically very complex, it is important to try to identify areas of focus on the genome with which to inform drug target research. The robust effects of neuroleptics on MDD symptoms is well-established, and it may be that certain (genetic) classes of individuals respond more or less to treatment based on common variation in the human genome. Because MDD is likely quite heterogeneous, it is important to try to identify genetic classes of cases that differ phenotypically. This view is supported by many labs currently attempting to genetically stratify severe psychopathology (13). Recent helpful research by Howard and colleagues has successfully replicated a genetic classification of MDD using PRS of ~20 medical and psychiatric phenotypes. Future research would benefit from examination of whether high-risk PRS classes are phenotypically distinct (whether clinical features differ) and also whether they are distinct with respect to risk within drug-relevant molecular pathways.

Were there enough non-EUR GWAS to complete a similar stratification analysis in CONVERGE, we would have attempted this in conjunction with our analysis of molecular pathway-based genetic MDD risk. PRS for >200 phenotypes can currently be calculated for EUR samples based on EUR GWAS, but Unfortunately the field lacks sufficient non-EUR samples for this type of analysis in CONVERGE. We hope that future GWAS of non-EUR samples will facilitate additional meaningful insights.

Importantly, these findings require replication and may not be generalizable to other ancestry groups or to male MDD cases and controls. While ascertainment in this study successfully minimized genetic heterogeneity due to sex differences and ancestry, this also limits the generalizability of GO:17144 drug metabolism pathway findings. Because the study rigorously ascertained recurrent MDD, and not single episode, it is also possible that the effects observed here are limited to severe and/or recurrent MDD.

Finally, there are problems inherent to any study of gene pathways. These include the field’s very limited molecular understanding of gene pathways and related sequelae, incompleteness of pathway databases, conflicting opinions about the independence/interdependence of gene pathways, and the elevated polygenicity of all psychiatric traits. To the first three points, we assert that it is studies like this that further develop the pathway literature and inform pathway database efforts. To the fourth point, it is critical for studies to account for genome-wide polygenic risk when testing for any signal relevant to a small portion of the genome. These points underscore the importance of careful analysis and independent replication.

## Acknowledgements

The CONVERGE Consortium gratefully acknowledges the support of all partners in hospitals across China, without whom this research would not be possible. This work was supported by the Wellcome Trust (WT090532/Z/09/Z, WT083573/Z/07/Z, WT089269/Z/09/Z) and by National Institute of Mental Health grant MH100549. The Wellcome Trust had no further role in study design, data collection, analysis, or interpretation, or in the writing of this report. This project was supported by NIH grants K01MH093731 (Dr. Docherty), KMH109765 (Dr. Adkins), MH100560 (Dr. Bacanu), AA022717 (Dr. Bacanu), AA021399 (Dr. Edwards), by a NARSAD Young Investigator Award (Dr. Docherty), and by the University of Utah EDGE Scholar Program.

